# Identification of radiation responsive proteins from Spleen and Intestine tissue of mice using Mass Spectrometry approach

**DOI:** 10.1101/2020.02.24.962647

**Authors:** Md. Shabir Husain, Faisal Ahmed, Ram Kumar, Akhilesh K Saini, Smriti Shubham, Saggere M Shasthry, Mohammad Tariq Salman

## Abstract

**Purpose:** Exposure to ionizing radiation (IR) can cause tissue damage, which is difficult to diagnose and treat as no biomarker is available for detection. We aimed to identify proteomic signature of radiation exposure (9.5Gry) in mice and to assess the utility of *Podophyllotoxin* extract (PTOX) in preventing radiation injury.

**Materials and Methods:** Spleen and Small intestinal (SI) tissues were taken from control and lethally irradiated mice at different time intervals with or without pre-treatment with *Podophyllotoxin* extract. Proteins were identified using Mass Spectrometry and matched with Peptide Mass Fingerprinting.

**Results:** We found multiple differentially expressed radiation responsive proteins from Spleen and SI tissues in irradiated mice at 24 hours and 30 days in comparison to healthy controls (p<0.05). Differentially expressed proteins like Chromosome transmission fidelity factor ano thath 18 homolog (CTF18) and Rho GTPase-activating protein from spleen and Acta_Mouse protein from SI were identified. These proteins disappeared at 48 hrs. after IR, but re-appeared after 13 days and fully recovered at 30 days in *Podophyllotoxin* treated group.

**Conclusions:** Such proteins may be useful in early detection of radiation exposure. Pre-treatment with *Podophyllotoxin* leads to recovery of the disappeared proteins and improved survival following exposure to irradiation.

## 1. Introduction

Gamma rays, a type of ionizing radiation (IR) can cause tissue damage by targeting macromolecules like DNA, proteins etc. (Hussain et al. 2017) (Lahtz et al. 2012),(Miura et al. 2007). Radiation exposure can happen either from the cancer therapies or from accidental exposure (nuclear plants etc.) with potential health hazards. The fear of IR related health hazards has necessitated the need for development of diagnostic tests, including identification of biomarkers for early diagnosis and treatment. The biomarkers might also be helpful for the development of radio protective agents to alleviate side effects of IR (Stochaj et al. 2006). There is limited literature on the diagnostic biomarkers and treatment options for radiation related injuries. So, we attempted to identify radiation responsive proteins using proteomics approach. Additionally, we also used *Podophyllotoxin (PTOX)* (from the Rhizome of <u>*P.hexandrum*</u>) prior to IR as it had shown some radio protective properties in our previous study (Ritorto and Borlak 2008), (Shih et al. 2015), Hussain et al 2017). *Podphyllotoxin* is an active cytotoxic natural product used as starting compound for the synthesis of anticancer agent, etoposide and teniposide (Ardalani H et al 2017), (Cai SJ et al 2019). Our present study focuses on proteomic analysis and identification of differentially expressed proteins in radiosensitive organs after exposure to lethal dose of IR and as well the utility of *P. Hexendram* as a radio protective agent.

## 2. Material and Methods

We used Two-Dimensional Gel Electrophoresis (2-DE), (Bio-Rad) and silver staining to separate and visualize proteins in 2-DE PAGE (polyacrylamide gel electrophoresis). Mass Spectrometry (MS) (Applied Bio systems 5800) was used to identify the differentially expressed proteins. A series of methods such 2-DE, Matrix-assisted laser desorption/ionization time of flight/ MS (MALDI-TOF-MS) and PMF based on Swiss-Port database searching were used to separate and identify differentially expressed proteins in gamma-ray irradiated mice model. Our study was performed on spleen and SI tissues taken from HC, vehicle control (VC), lethally irradiated and PTOX + lethally irradiated group (female, 6 - 8 weeks old strain ‘A’ mice). One of the group only irradiated, this group of mice were death after 13^th^ days, we were not collected the tissues samples. Excised tissues were subjected to 2-DE to generate differential proteome data after 24hr and 48hr of irradiation. In another set of mice, the expression of proteins was studied in *Podophyllotoxin* pretreated mice 24hr, 48hr, 13days and 30days after irradiation. The drugs 1:2 ratio of rutin from purified *Podophyllotoxin.* We have checked the chemical toxicity and purity by High Performance Liquid Chromatography (HPLC). *PTOX* (6.5 mg /kg) was administered intramuscularly 60 minutes prior to irradiation. (Table.1). Differentially expressed proteins spots were selected in spleen by PD Quest analysis from 2-DE gels of irradiated samples with comparison to Controls.

**Table: 1.** Groups of mice: 3 Sections of mice in 9 different Groups: Section ‘A’ Healthy Control (HC), Section ‘B’ Radiation (R), Section ‘C’ PTOX with Radiation (PTOX+R). And two mice of vehicle control (VC) in group III.

| Section | Section 'A' |  |  | Section 'B' |  |  |  | Section 'C' |  |  |  |
| --- | --- | --- | --- | --- | --- | --- | --- | --- | --- | --- | --- |
|  | Healthy Control (HC) |  |  | Radiation (R) |  |  |  | *Podophyllotoxin* (PTOX) with Radiation (PTOX+R) |  |  |  |
| Groups | Group I | Group II | Group III | Group IV | Group V | Group VI | Group VII | Group VII | Group IX |  |  |
| Samples | HC<br>No PTOX or Radiation | D<br>Only PTOX | VC<br>Vehicle Control | HC<br>No PTOX or Radiation | R 24h<br>Only radiation | R 48h<br>Only radiation | Only radiation<br>Mice date after 13<sup>th</sup> days | PTOX+R<br>24 h | PTOX +R<br>48h | PTOX +R<br>13 days | PTOX+R<br>30days |
| No. of Mice | 3 | 3 | 2 | 3 | 3 | 3 | 3 | 3 | 3 | 3 | 3 |

### 2.1. Mice

Strain “A” female mice (6-8 weeks old) were obtained from Radio-protective Drug Development Group, Division of Radiation Biosciences, Institute of Nuclear Medicine and Allied Sciences (INMAS), Delhi, India for the radiation experiments. Mice were acclimatized to the animal care facility in institutional for the experiment. They were accommodated 12:12 light: dark cycle under 22 °C and humidity was (40– 60%), and illumination. We adhered to the Institutional Animal Care facility and Committee (IACUC) guidelines for maintenance and use animals.

#### 2.2. a. Group of mice with irradiation

Nine groups comprising 3 mice in each, were irradiated with 9.5 Gray gamma radiations and sacrificed for collection of spleen and SI tissue samples at different time intervals as mentioned in the table-1, after being anesthetized by intra peritoneal Pentobarbital Sodium (100 mg /Kg).

#### b. Drugs and chemicals

Purified *Podophyllum hexandrum* was given at a dose of 6.5 mg/kg, intra muscular, i.m., 60 minutes prior to irradiation. The chemical formula of *Podophyllotoxin(PTOX)* is C_22_H_22_O_8_ with molecular weight of 414.41. PTOX belonged to vascular plants, it has been isolated from roots or rhizomes of plant (Ardalani H et al 2017).

### 2.3. Protein separation and identification

#### 2.3. a. Separation of proteins by 2-DE PAGE

We used Immobilized pH gradient (IPG) 17cm, pH 4–7 (Bio-Rad Laboratories, Richmond, CA) strips for rehydrated of 70μg of spleen and SI proteins extracts (24hr, 48hr, D+R 24hr, D+R 48hr, D+R 13days, D+R 30 days with HC, irradiated sample and sham irradiated control) for 12hr at normal temperature. In IPG apparatus (Bio-Rad Laboratories, Richmond, CA) focused for 8hr for a total of 40,000 V-hr. Focused IPG strips were then equilibrated in the presence of 1% DTT and subsequently with 2.5% iodoacetamidethe containing 50 mM Tris-HCl buffer, pH 8.8, 6 M urea, 20% v/v glycerol, 0.1% w/v SDS for the equilibration buffer and equilibrated for 20 minutes each with shaking. To remove the buffer, strips were then loaded for 12.5% SDS-PAGE gel on 100 V/ 1 hrs. We did the tests in triplicates (of 64 SDS gels) for the identifying proteins.

#### 2.3. b. Identification of Proteins by high throughput Mass Spectrometry

We used the high-throughput and quantitative MALDI-TOF/MS for protein structure, function, modification and global protein dynamics. Identified novel proteins, peptides of amino acid and PMF to the basis of mass and charge (m/z) ratios which were mentioned in tables -2.a., 2.b and 2.c.

**Table: 2. A.**
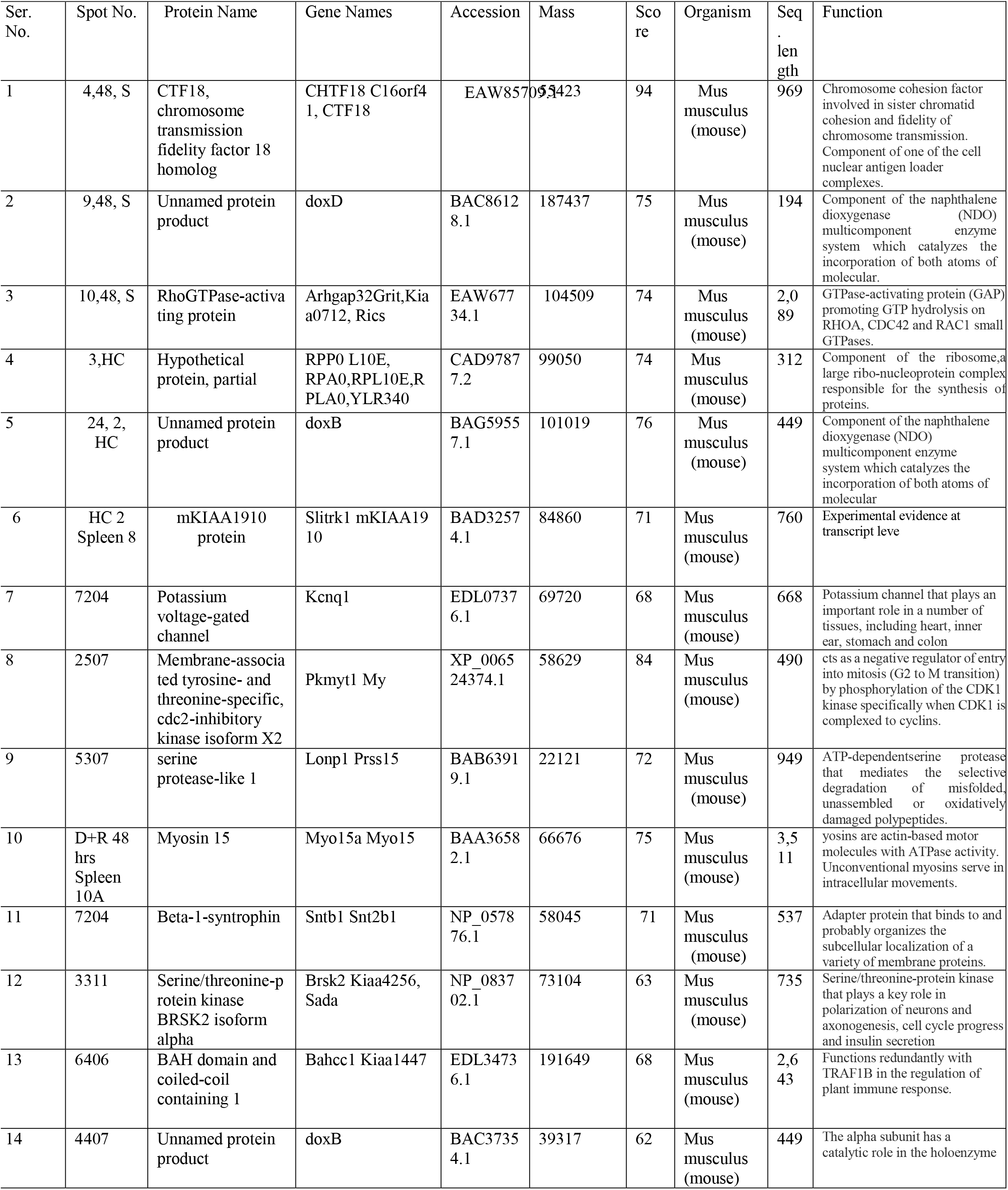
List of identified differentially expressed proteins by MALDI-TOF/ Mass Spectrometry at the different time irradiated mice Spleen tissues with sham healthy control (HC).

**Table. 2, B:**
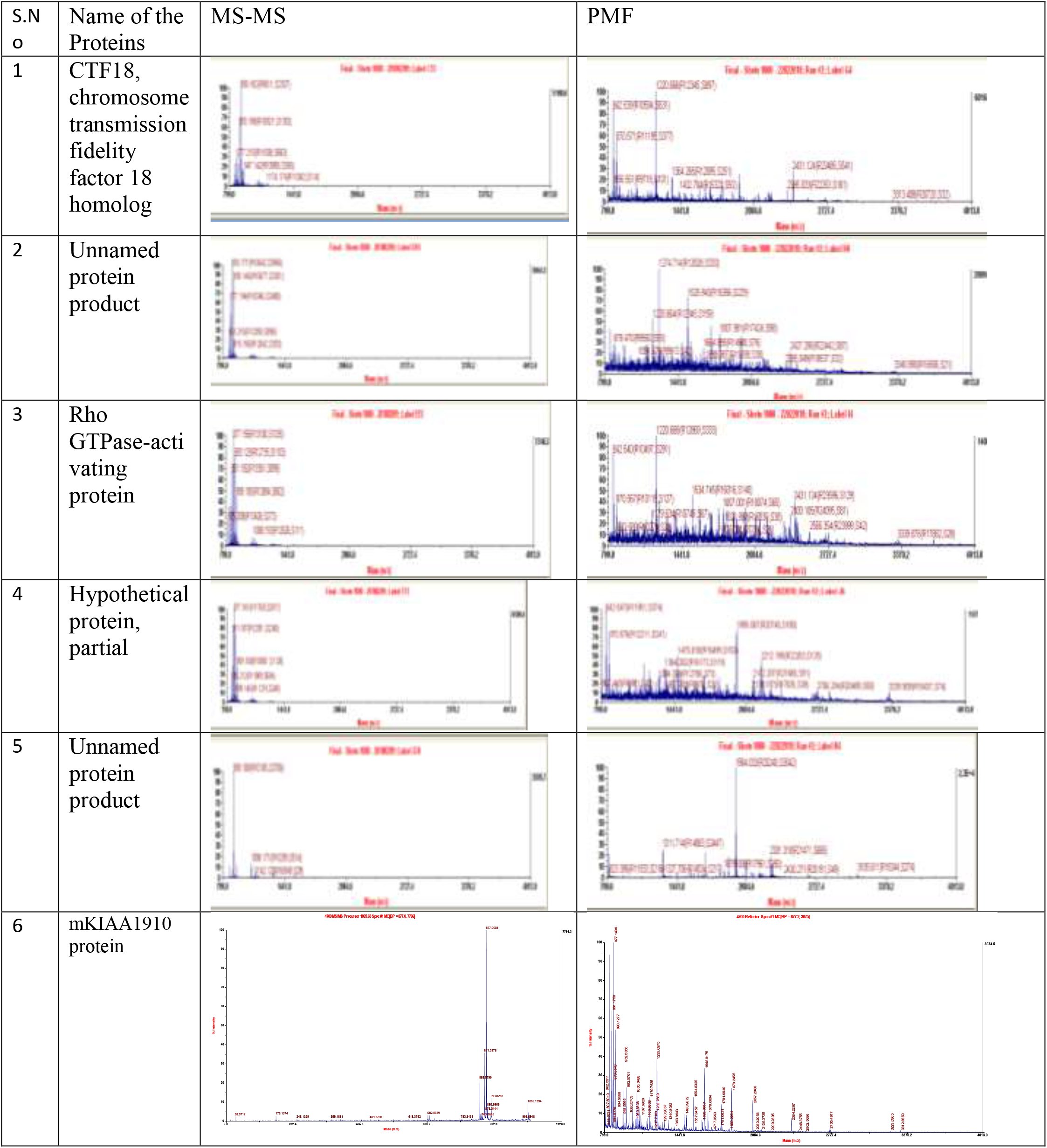
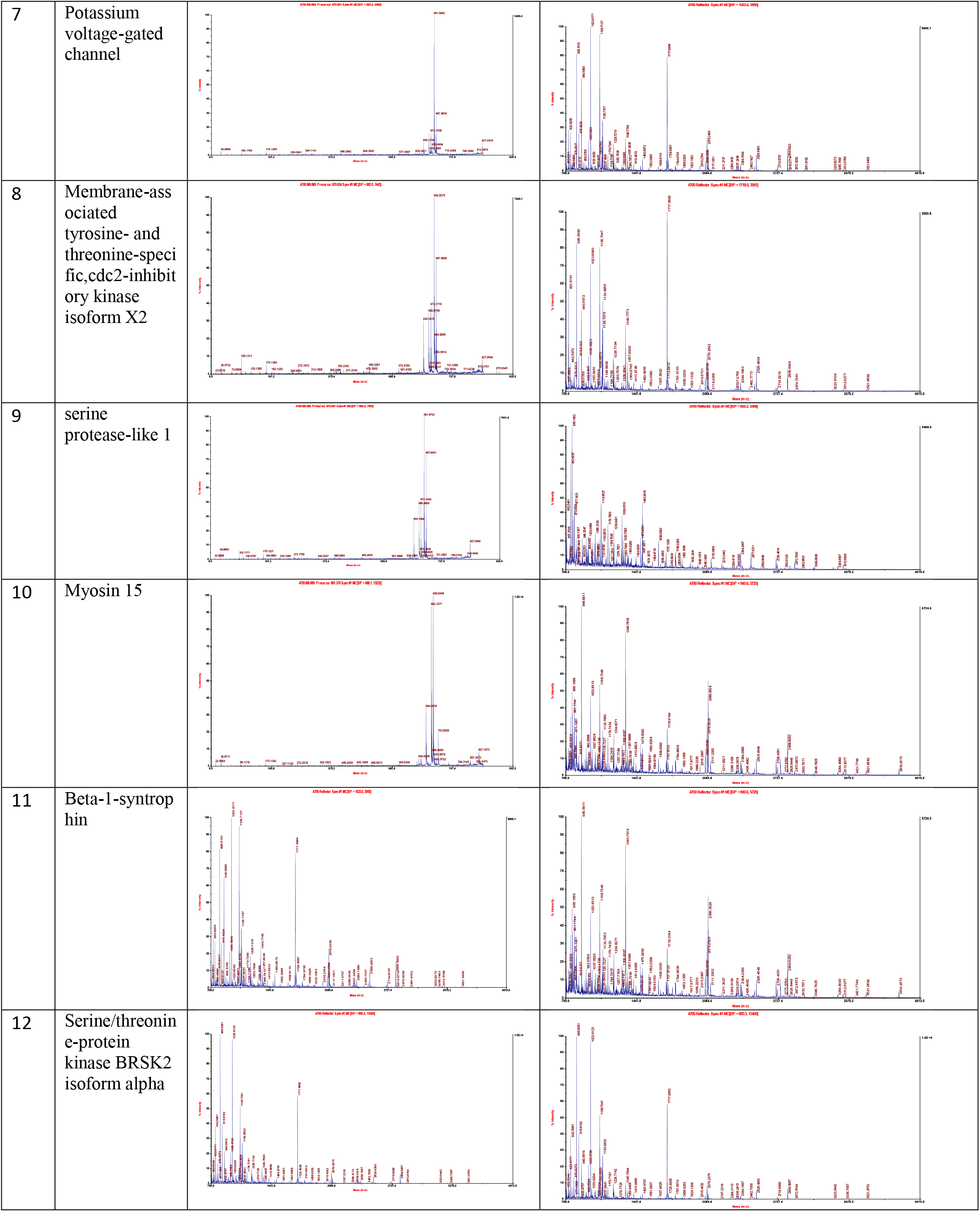
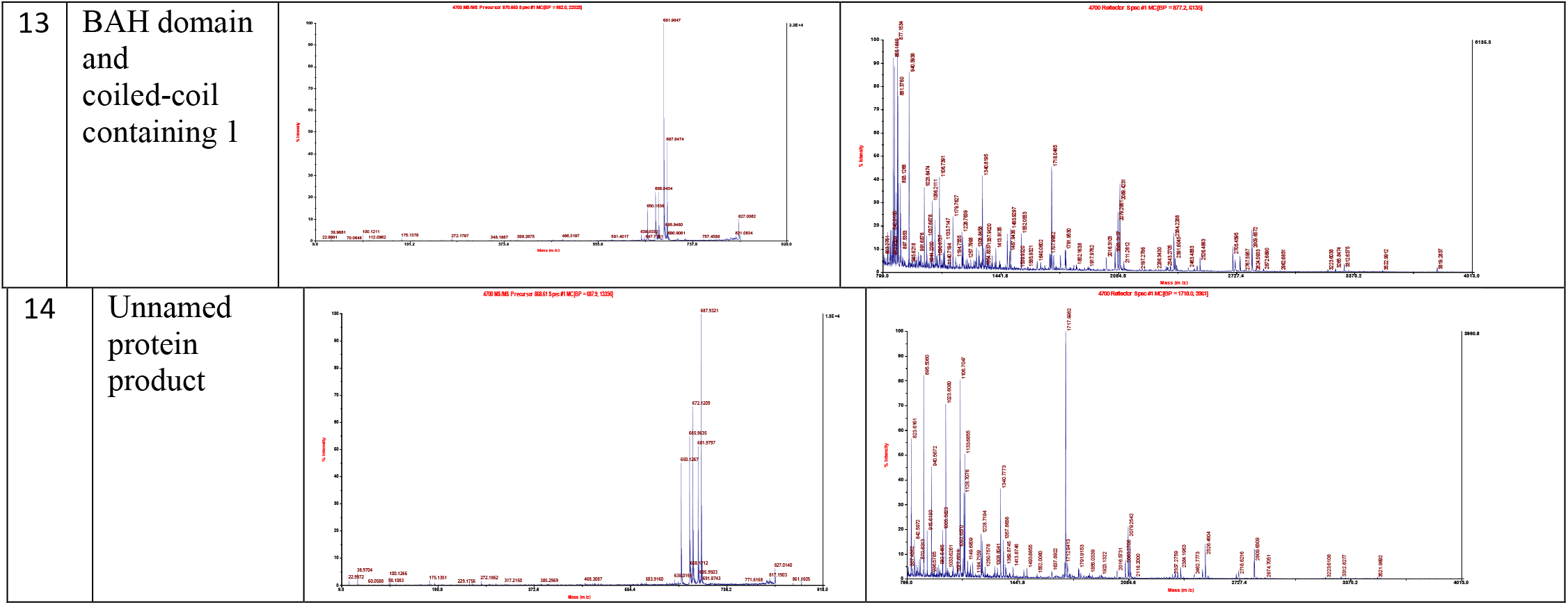
List of MS-MS and PMF spectra of the amino acid from identified proteins from spleen by Mass Spectrometry.

**Table: 2. A.**
| S.N<br>o | Name of the<br>Proteins | MS-MS | PMF |
| --- | --- | --- | --- |
| 1 | CTF18,<br>chromosome<br>transmission<br>fidelity<br>factor 18<br>homolog |  |  |
| 2 | Unnamed<br>protein<br>product |  |  |
| 3 | Rho<br>GTPase-acti<br>vating<br>protein |  |  |
| 4 | Hypothetical<br>protein,<br>partial |  |  |
| 5 | Unnamed<br>protein<br>product |  |  |
| 6 | mKIAA1910<br>protein |  |  |
| 7 | Potassium voltage-gated channel |  |  |
| 8 | Membrane-associated tyrosine- and threonine-specific,cdc2-inhibitory kinase isoform X2 |  |  |
| 9 | serine protease-like 1 |  |  |
| 10 | Myosin 15 |  |  |
| 11 | Beta-1-syntrophin |  |  |
| 12 | Serine/threonine-protein kinase BRSK2 isoform alpha |  |  |

|  |  |
| --- | --- |
| 13 | BAH domain and coiled-coil containing 1 |
| 14 | Unnamed protein product |

**Table: 2, C.** Identification of differentially expressed protein from SI tissues by MALDI-TOF/ Mass Spectrometry.

**Table. 2, B:**
| Spot No. | Protein Name | Gene Names | Accession | Mass | Score | Organism | Seq. length | Function |
| --- | --- | --- | --- | --- | --- | --- | --- | --- |
| HC SI | T-cell receptor alpha chain precursor/ T-cell surface glycoprotein CD3 zeta chain | Cd247 | AAT2744.1 | 630141 | 49 | Mus musculus (mouse) | 164 | Part of the TCR-CD3 complex present on T-lymphocyte cell surface that plays an essential role in adaptive immune response. |
| 48 R SI | Acta_Mouse | Acta2 | 42381 | 87909 | 93 | Mus musculus (mouse) | 376 | Actins are highly conserved proteins that are involved in various types of cell motility and are ubiquitously expressed in all eukaryotic cells. |

### 2.4. Analysis of data

We scanned all 2-DE gel by 16 bit scale and images in TIF form, Bio-Rad Laboratories, Richmond, CA. scanner and digitized image data was analyzed using PDQuest software (Bio-Rad Laboratories, Richmond, CA). Protein, and identified peptides matching between all the PMF were viewed and normalized by the relative intensity of individual proteins in spleen and SI tissues samples.

### 2.5. Trypsin digestion and extraction of peptides

We cut the proteins spots from gel and digested by trypsin enzyme (Shevchenko et al. 1996). Digested proteins were placed in 10mM NH_4_HCO_3_ for 5min and in equal volume of ACN for next 15min. Then, mixed trypsin enzyme in digested proteins samples, the concentration of trypsin was 20ng/μl (Mass Spectrometry grade, Promega), and placed the samples overnight at 37 °C for 12 to 15 hours. St Louis, MO, USA, thensonicated in water bath for 30 minutes to remove the ACN and stored at –20°C till further use for MS analysis (Biswas et al. 2013).

### 2.6. Identification of Proteins by MS and match the identified peptides in the MASCOT Database

The reconstituted peptide samples (1 μL) were analyzed by MS. The obtained peak lists were further searched using the available online MASCOT by matrixscience.com, then we used uniprot database for the important functions of the identified protein.

## 3. RESULTS

### 3.1. Identification of differentially expressed proteins by Mass Spectrometry

We found sixteen differentially expressed proteins from Spleen and eight from SI tissues of irradiated mice in comparison with HC. The 2-DE SDS gels were analyzed by the PD Quest and proteins were identified by high throughput technique MS. We found differences between HC and irradiated mice at 24 hours and onwards, till 30 days (p<0.05) with loss of the identified proteins. These proteins started to reappear in mice treated with G-003+R, by day 13 and nearly normalized by day-30 (p < 0.0001 in comparison to 24 hrs.) (Figure: 1). The protein Acta-mouse was identified from SI tissues at different time period of irradiation and Drug with radiation (Figure. 2), name and function of the identified proteins are mentioned in Table 2. A, The proteins such as Chromosome transmission fidelity factor 18 homolog (CTF18), Unnamed protein product, Rho GTPase-activating protein, Hypothetical protein partial, etc. were from Spleen tissues. They were radiation responsive proteins which disappeared at 48 hrs. IR re-appeared after 13 days and fully recovered at 30 days.

**Figure: 1.**
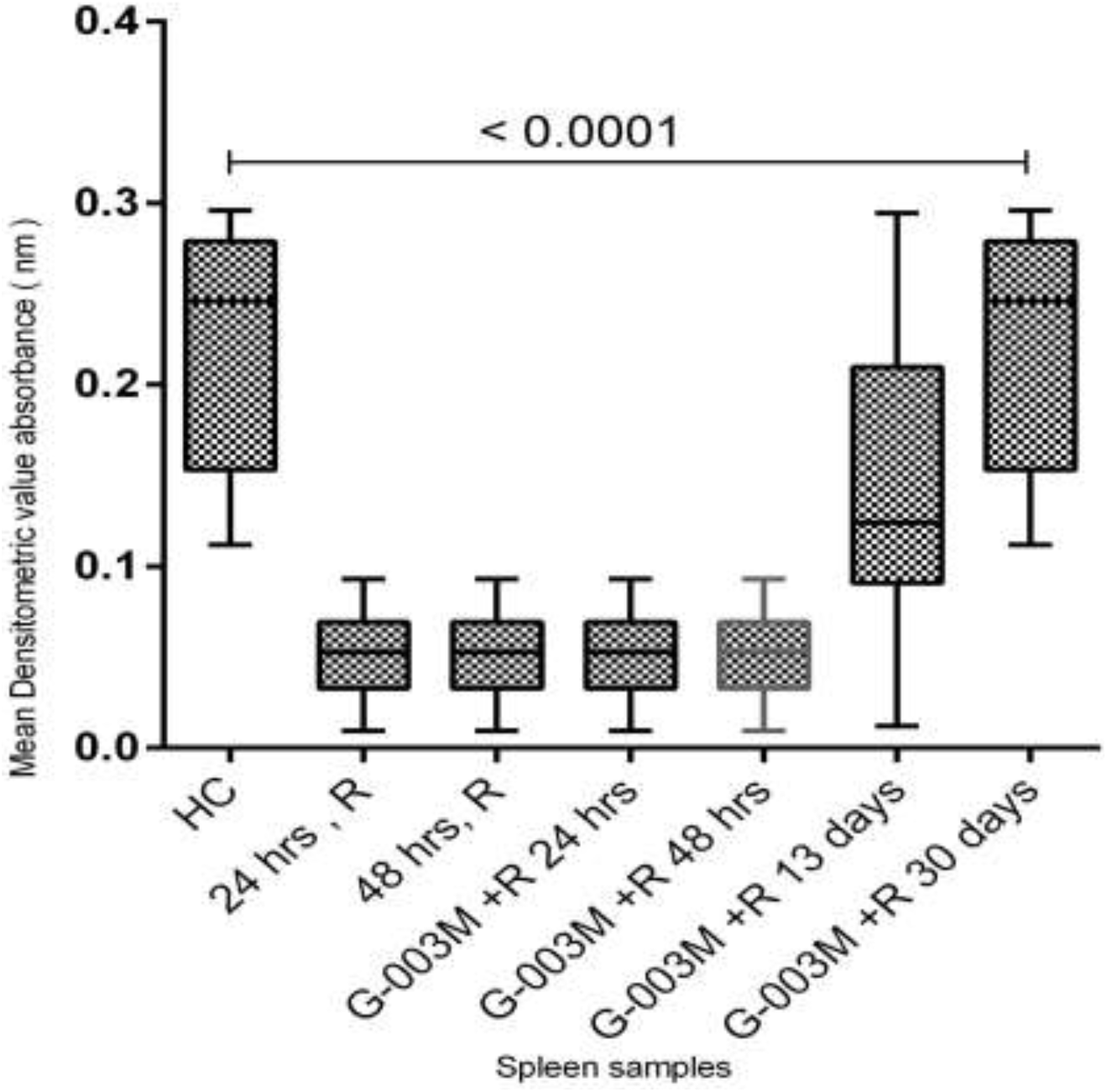
The status of most differently expressed proteins from spleen tissues of mice, at the different time intervals of the IR and *Podophyllotoxin* (PTOX), HC, PTOX +R 13days and PTOX +R at 30 days.

**Figure:2.**
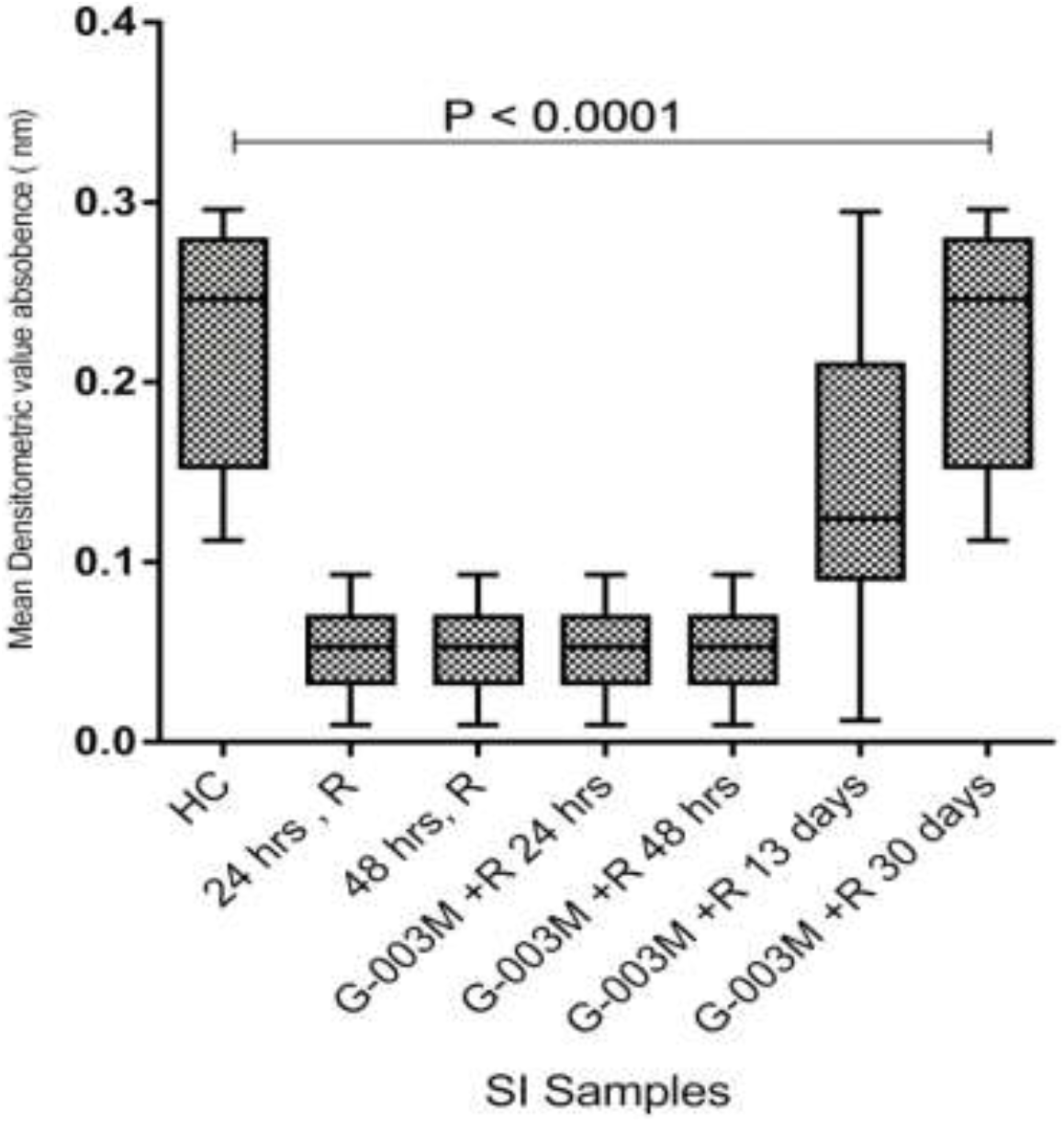
The status of most differently expressed proteins from SI tissue of mice, at the different time intervals of the IR and *Podophyllotoxin* (PTOX), HC, PTOX +R 13days and PTOX +R at 30 days.

### 3.2. MS/MS and PMF analysis of splenic tissue of radiated mice (G003M +radiation group) in comparison to healthy control and at different time periods

All differentially expressed down regulated proteins were identified (Table: 2.B.) from 24 hrs. and 48 hrs. G-003M+irradiation group, by MS/MS analysis. The identified proteins were confirmed by Mascot database search engine. These proteins were down regulated at 24 hrs. after radiation exposure in comparison to control and started to reappear after day-13 and recovered completely by day-30 in the mice receiving radiation and pretreated with G-003M. CTF18 (chromosome transmission fidelity factor 18 homolog) Unnamed protein product, RhoGTPase-activating protein, Hypothetical protein, partial, mKIAA1910 protein, serine protease-like 1 etc.

### 3.3. MS/MS and PMF analysis of SI tissues of radiated mice (G003M +radiation group) in comparison to healthy control at different time periods

We identified two important proteins, T-cell receptor alpha chain precursor/ T-cell surface glycoprotein CD3 zeta chain from HC group and Acta Mouse from 48 hrs irradiation from SI tissue. Name and associated function of all The radiation-responsive proteins (mentioned in Table: 2. C and D) from SI disappeared by 48 hrs after the IR, re-appeared after 13 days and fully recovered at 30 days. The MS/MS Peptides of all the identified proteins matched with PMF spectra and correlated with Protein sequence coverage in validation. 2-DE gels images from SI tissue were also compared between HC and G-003M + irradiation (in different groups). The details of the identified proteins and their important functions have been mentioned in the table: 2 C, D).

**Table: 2, D.**
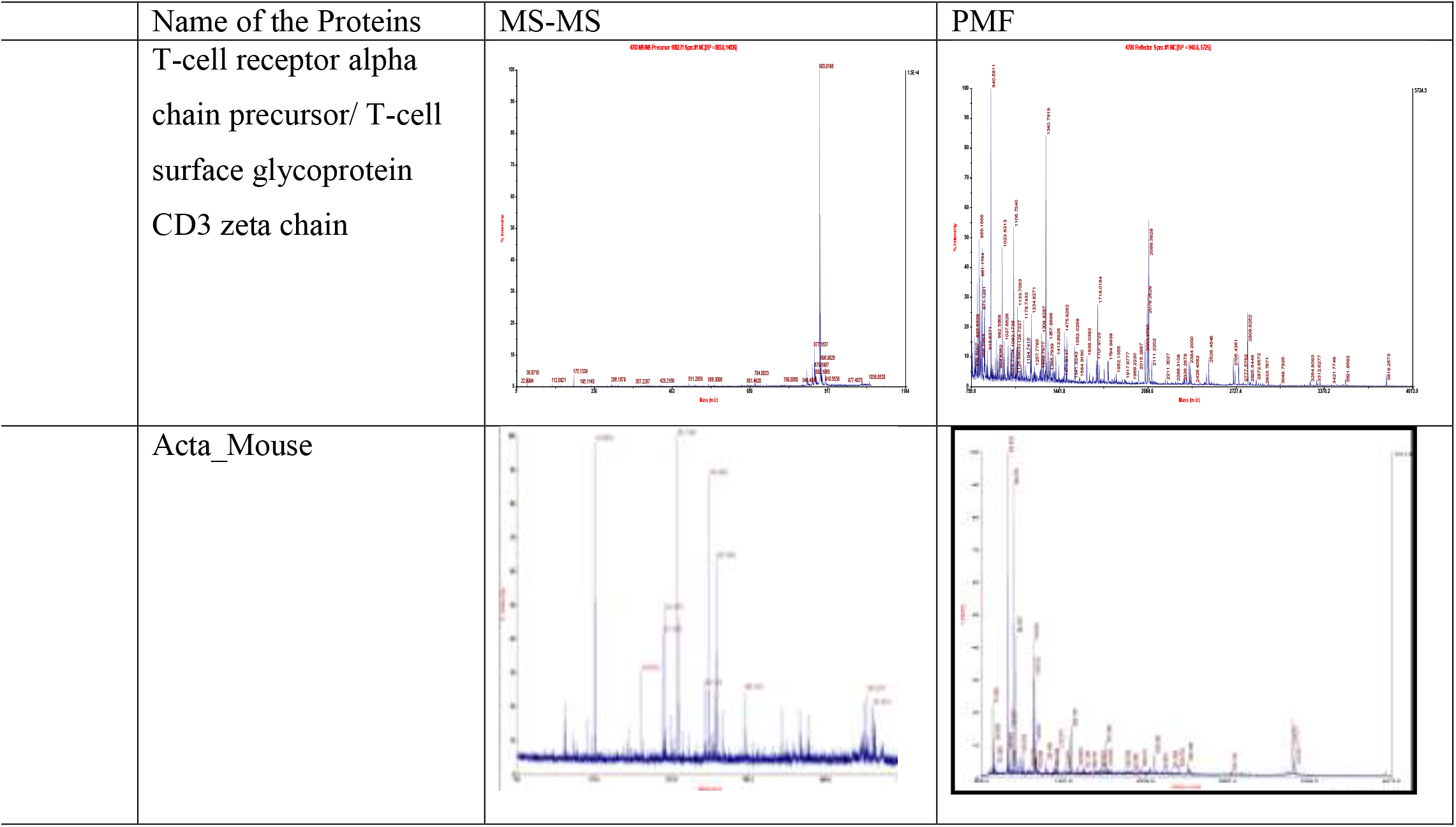
MS-MS and PMF spectra of identified Proteins from SI tissue samples.

**Table: 2, C.**
|  | Name of the Proteins | MS-MS | PMF |
| --- | --- | --- | --- |
|  | T-cell receptor alpha chain precursor/ T-cell surface glycoprotein CD3 zeta chain |  |  |
|  | Acta_Mouse |  |  |

## 4. DISCUSSION

We identified radiation responsive differentially expressed proteins after lethal dose radiation exposure from the Spleen and SI tissues of mice, which are specifically down regulated after radiation exposure and reappear later with recovery. The identified proteins (I)Chromosome transmission factor 18 homolog, it is sister chromatid and involved in transmission of the chromosomes- CTF8, DCC1, RFC2, RFC3, RFC4 and RFC5 and also with replication of DNA. (II)Unnamed protein product- is involved in enzymatic system which catalyzes molecular oxygen atoms such as dihydroxy-1, 2-dihydronaphthalene and may thus be involved in the reactive oxygen species generation post radiation exposure. (III) GTPase-activating protein (GAP), a ribosomal Protein which promotes the reaction of hydrolysis, involved in regulation of cell growth and pathways of the cell signaling (Qin et al. 2015). (IV)Unnamed protein product- involved in the amino acid sequencing. (V) mKIAA1910 protein- a potassium channel. We additionally identified 14 significant radioresponsive proteins, such as, Myosins (involved with cellular movement), Beta-1-syntrophin (membrane protein linking to many subcellular functions including cell cyclr progression etc.), Unnamed protein **product,** (suppresses cell growth, differentiation and angiogenesis in numerous types of carcinoma by interacting with certain cell cycle proteins, with CDK2 and DNA polymerase). Further studies are required to assess the relevance and exact roles of the above identified radiation responsive proteins and the mechanism of the action of the radio protective drug, the PTOX. Mice survived despite exposure to lethal dose of radiation (9.5Gy) because of prior treatment with *PTOX* and the radiosensitive down-regulated proteins normalized by 30 days. This indicated that the proteins could be prospective biomarkers for radiation exposure. Identification of down regulation of these proteins could help in identification and early treatment of patients after radiation exposure and recovery of such proteins could serve as biomarker of recovery. A battery of biomarkers identified by comparative proteomics approaches in various tissues could represent a promising and potent tool for the discovery of innovative radiation biomarkers which could be helpful in better management of radiation related health hazards.

## Abbreviations

IR: ionizing radiation
SI: Small intestinal
2-DE: Two-Dimensional Gel Electrophoresis
IPG: Immobilized pH gradient
MALDI-TOF/ MS: Matrix-Assisted Laser Desorption Ionization -Time of Flight /Mass Spectrometry
Podophyllotoxin (PTOX) +R or PTOX + R: with radiation
hrs: hours
Gy: Gray

## ACKNOWLEDGEMENTS

We would like to acknowledge Defence Research & Development Organization (DRDO), Ministry of Defence for providing financial support to carry out the research work. Dr. Manju Gupta, Dr. Ajaswrata Dutta,INMAS, DRDO, and Dr. Sagarika Biswas, CSIR-IGIB, New Delhi for help in experiment.

## Notes

**Financial Support:** This work is funded by the Defense Research and Development Organization (DRDO), India.

**Disclosure:** All authors have declared no conflict of interest.

## REFERENCES

Hussain S, Dutta A, Sarkar A, Singh A, Gupta ML, Biswas S. 2017. Proteomic analysis of irradiated lung tissue of mice using gel-based proteomic approach. Int J Radiat Biol. 93(4):373–380. eng.

Lahtz C, Bates SE, Jiang Y, Li AX, Wu X, Hahn MA, Pfeifer GP. 2012. Gamma irradiation does not induce detectable changes in DNA methylation directly following exposure of human cells. PLoS One. 7(9):e44858. eng.

Miura Y, Kano M, Yamada M, Nishine T, Urano S, Suzuki S, Endo T, Toda T. 2007. Proteomic study on X-irradiation-responsive proteins and ageing: search for responsible proteins for radiation adaptive response. J Biochem. 142(2):145–155. eng.

Qin J, Rajaratnam R, Feng L, Salami J, Barber-Rotenberg JS, Domsic J, Reyes-Uribe P, Liu H, Dang W, Berger SL et al. 2015. Development of organometallic S6K1 inhibitors. Journal of medicinal chemistry. 58(1):305–314.

Ritorto MS, Borlak J. 2008. A simple and reliable protocol for mouse serum proteome profiling studies by use of two-dimensional electrophoresis and MALDI TOF/TOF mass spectrometry. Proteome Sci. 6:25. eng.

Shevchenko A, Jensen ON, Podtelejnikov AV, Sagliocco F, Wilm M, Vorm O, Mortensen P, Boucherie H, Mann M. 1996. Linking genome and proteome by mass spectrometry: large-scale identification of yeast proteins from two dimensional gels. Proc Natl Acad Sci U S A. 93(25):14440–14445. eng.

Shih CJ, Chen SC, Weng CY, Lai MC, Yang YL. 2015. Rapid identification of haloarchaea and methanoarchaea using the matrix assisted laser desorption/ionization time-of-flight mass spectrometry. Sci Rep. 5:16326. eng.

Stochaj WR, Berkelman T, Laird N. 2006. Preparative 2D Gel Electrophoresis with Immobilized pH Gradients: IPG Strip Equilibration. CSH Protoc. 2006(5). eng.

Hamidreza Ardalani, Amir Avan, Majid Ghayour-MobarhanPodophyllotoxin: a novel potential natural anticancer agent 2017.

Cai SJ, Liu Y, Han S, Yang C.Brusatol, an NRF2 inhibitor for future cancer therapeutic2019.

